# Morphological evolution of bird wings follows a mechanical sensitivity gradient determined by the aerodynamics of flapping flight

**DOI:** 10.1101/2022.09.23.509279

**Authors:** Jonathan Rader, Tyson L. Hedrick

## Abstract

The physical principles that govern the function of biological structures also mediate their evolution, but the evolutionary drivers of morphological traits within complex structures can be difficult to predict. We used morphological traits measured from 1096 3-dimensional bird wing scans from 178 species to test how two frameworks for relating morphology to evolution interact. We examined whether the modular organization of the wing into handwing and armwing regions, or the relationship between trait morphology and functional output (i.e. mechanical sensitivity, driven here by flapping flight aerodynamics) dominate evolutionary rate (*σ*^*2*^) and mode. Our results support discretization of the armwing and handwing as morphological modules, but morphological disparity and *σ*^*2*^ varied continuously with the mechanical sensitivity gradient and were not modular. Thus, mechanical sensitivity should be considered an independent driver of evolutionary dynamics, distinct from morphological modularity.

**Teaser:** Mechanical sensitivity drives wing shape evolution in birds and may be fundamental to the evolution of biomechanical systems.

## Introduction

Form – function relationships are one of the pillars of biodiversity. Morphological features have diverged in size and shape among lineages and impart different abilities to interact with the environment and compete for finite resources (*1*–*4*). The evolution of morphological traits is not always directly and solely linked to their function, though. Shared development and function can lead to coevolution of phenotypic traits (*5, 6*). Individual traits within a system can contribute in varying degrees to the functional output of the whole. Furthermore, the degree of integration among these coevolving traits and their organization into mosaics of semi-independent modules are mediated by their shared development and by the magnitude of their impact on functional output (*7, 8*). In a biomechanical context, the strength of relationships between morphological traits and their mechanical function (termed mechanical sensitivity) may be an important driver of their evolutionary dynamics (i.e. tempo and mode) (*9*–*11*).

The first description of the relationship between mechanical sensitivity and evolutionary dynamics focused on four-bar linkage systems, particularly in the jaws of teleost fish (*12, 13*) and the raptorial appendages of mantis shrimp (*14*). Here, each link can be thought of as a discrete morphological module. The modules with the greatest impact on the transmission of force or motion in a four-bar linkage system also have the greatest mechanical sensitivity (*9*), which correlates with a shift in evolutionary mode (from Brownian motion toward Ornstein-Uhlenbeck) and to a higher evolutionary tempo (*10, 11*). However, despite finding a similar coupling of mechanical sensitivity and evolutionary dynamics in the four-bar linkage systems in two disparate taxa, the generalizability of these results remains hampered by a lack of comparable studies of other morphological traits in biomechanical systems beyond the four-bar linkage (*10*). As such, studying morphological modularity and evolution in systems with different biophysical interactions, such as the fluid-structure interactions in flight or swimming, can fill some of the missing picture of the patterns and processes that shape evolution of complex biological structures. We investigated how evolutionary dynamics of wing shape in birds have responded to the interplay between the mechanics of aerodynamic force production in flight and morphological modularity within the wing.

### Form begets function in bird wings

Birds are diverse in their ecology and behavior, which manifests in differences in their flight style, morphology (see Fig. 1a), and performance. Bird wings must produce lift to support body weight during flight and asymmetrical forces for maneuvering. They must function at cruising speed, at low airspeeds during landing and maneuvering flight, and during supra-normal efforts for pursuit or escape flight. Furthermore, bird wings may experience trade-offs and constraints imposed by their structure and evolutionary development.

**Figure 1.**
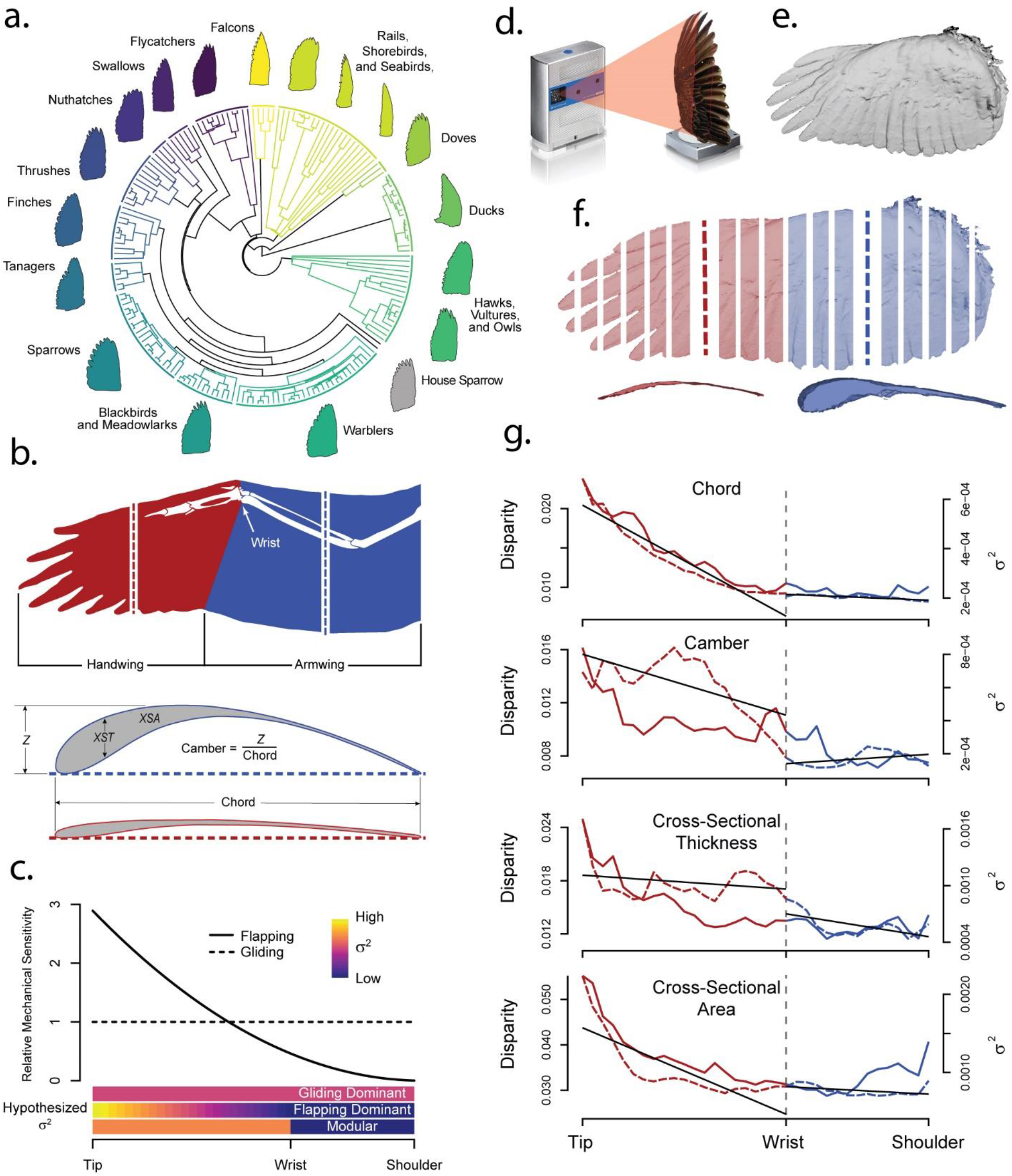
Bird wings are complex biological structures with phylogenetically structured morphological variability (a.) and are composed of musculoskeletal and integumentary elements (b.). Flight feathers form the aerodynamic surface of the wing, which is divided into two regions separated by the wrist, the handwing and the armwing (b.). Gliding flight imposes consistent mechanical sensitivity along the wing span, (c., dashed line) Aerodynamic forces and inertial moment increase as the square of the distance from the base of the wing during flapping flight ^40^, creating an alternative model for wing mechanical sensitivity (c., solid line). The evolutionary signature of this mechanical sensitivity gradient might be discretized by modularity in the wing (c., *σ*2). We used a laser scanner (d.) to capture surface scans (e.) of 178 species of birds (a.). We divided the wing into chord-wise slices (f.) along the span, and measured chord, camber, cross-sectional thickness (*XST*), and cross-sectional area (*XSA*) from each slice (see pane b.). Morphological disparity (dashed line) and evolutionary rate (*σ*^*2*^, solid line) for all these shape traits were greater in the handwing (red) than in the armwing (blue), and especially so near the wingtip. Regression discontinuity analyses (RDA) of morphological disparity (solid black lines) showed significant discontinuity across the wrist joint (see Table 1). The distinction across the wrist was less clear for *σ*^*2*^, except in wing chord and marginally in *XST*.

The geometry of a wing influences how it interacts with the air, and thus the lift and drag forces that it generates (*15*). Consequently, wing shape in birds is related to flight and migration behavior (*16*–*21*) and numerous other aspects of avian biology (*22*–*26*). Much work has been done describing how planform (2-dimensional) wing shape is related to avian aerodynamics (e.g. *27*–*32*). However, wings are not two-dimensional structures. Three-dimensional (3D) shape attributes such as wing camber (the upward curvature of the wing’s surface, see Fig. 1) contribute to aerodynamic forces produced by the wing (e.g., *33, 34*), and the distribution of mass along the wing span impacts the cost of flapping (*35*) and maneuverability (*36, 37*). Because 3D attributes of the wing are tied to its function, they are also potentially evolutionarily labile and tunable features, worthy of consideration in the story of avian wing evolution (*34, 19*).

**Table 1.** Phylogenetic signal, morphological disparity, evolutionary tempo (*σ*^*2*^) and regression discontinuity analyses for four wing shape traits: wing camber, chord, cross-sectional thickness (*XST*), and cross-sectional area (*XSA*).

| Shape Trait | Mean Phylogenetic Signal<br>(HW $\lambda, K$ \| AW $\lambda, K$) | | Mean Disparity<br>(HW \| AW) | | Mean $\sigma^2$<br>(HW \| AW) | | Disparity RDA<br>effect $p$ | $\sigma^2$ RDA<br>effect $p$ |
| --- | --- | --- | --- | --- | --- | --- | --- | --- |
| Camber | **0.69, 0.25** | **0.71, 0.28** | **0.014** | 0.008 | **0.0005** | 0.0003 | **0.000156\*** | 0.721920 |
| Chord | **0.86, 0.32** | **0.90, 0.41** | **0.012** | 0.008 | **0.0004** | 0.0002 | **0.000114\*** | **0.000351\*** |
| $XST$ | **0.64, 0.17** | **0.67, 0.16** | **0.018** | 0.013 | **0.0009** | 0.0006 | **0.0257\*** | **0.0282\*** |
| $XSA$ | **0.79, 0.19** | **0.76, 0.21** | **0.035** | 0.030 | **0.0012** | 0.0009 | **0.0141\*** | 0.487 |

\* $P < 0.05$

### Are birds wings modular?

Though the wing feathers create a generally contiguous wing surface, the avian wing is composed of multiple anatomical subunits (*38*). The most obvious of these are associated with the major skeletal regions of the forelimb (Fig. 1b). The portion of the wing associated with the radius and ulna (and to a lesser degree, the humerus) is the armwing (AW), and includes the bony elements, muscles, tendons, and the secondary and tertial portions of the feathered wing surface (*38*). The AW also supports the propatagium on its leading edge. The handwing (HW) is comprised by the bones of the wrist and hand as well as the primary portion of the feathered wing surface, but with minimal contribution from muscles and tendons (*38*). These two wing regions may be under differential selective pressures, or subject to different selective or developmental tradeoffs and constraints leading to evolutionary regionalization and modularity within the wing (*7, 38, 39*). For these reasons, we hypothesized that morphological modularity exists in the wing, dividing it into discrete armwing and handwing modules.

### Are evolutionary dynamics in the wing modular?

Inertial effects and aerodynamic forces from flapping flight increase as the square of the distance from the base of the wing, with the greatest effects at the tip of the wing (*40*). This gradient of aerodynamic and inertial effects produces a gradient of mechanical sensitivity along the length of the wing, smoothly crossing the hypothesized junction of handwing and armwing modules (Fig. 1; Appendix 1). If mechanical sensitivity is tied to the evolution of wing shape as it is in four-bar linkages (*11*), the gradient of mechanical sensitivity along the wing could result in a similar gradient of evolutionary dynamics of shape traits. We identified three idealized patterns that might characterize evolutionary dynamics in the wing: 1) they might follow the mechanical sensitivity of steady-state (gliding) flight and be uncorrelated with the force gradient imposed by flapping. In this case, there is no base-to-tip gradient of mechanical sensitivity, and no hypothesized differences in evolutionary tempo or mode along the wing regardless of modularity (see Fig. 1c). Alternatively, 2) evolutionary dynamics could follow the base to tip gradient in mechanical sensitivity established by flapping flight, irrespective of whether the wing is organized into morphological modules, producing a smooth root to tip pattern of increasing evolutionary tempo (Fig. 1c). 3) If flapping flight governs mechanical sensitivity but evolution acts upon modules within the wing, the HW region would have a distinctly faster evolutionary tempo and possibly a different evolutionary mode than the AW, but tempo would vary less within these modules than among them (Fig. 1c). Overall, the importance of flapping in avian flight combined with anatomical division of the wing into HW and AW regions led us to favor hypothesis 3, that shape traits would display greater morphological disparity near the tip of the wing, and that evolutionary tempo would also increase from the base of the wing toward its tip, with the differences exhibiting a regionally discontinuous pattern between the HW and AW. We investigated these hypotheses using 3D surface scans of wings from 178 species representing 15 major lineages of birds (Fig. 1a), providing a basis for exploring regionalization and modularity of avian wing morphology and evolution.

## Results

We scanned 1096 wings representing 178 species of birds with an average sample size of 6 individuals per species. Median wing camber across all slices, averaged within each species, ranged from 0.061 to 0.169, with an overall mean of 0.105. Armwing (AW) camber was greater than that in the handwing (HW, median ± MAD: 0.11 ± 0.021 vs. 0.072 ± 0.017). Mean chord ranged from 30.6 mm to 249.9 mm, with an overall mean of 64.3 mm. Median chord (scaled by dividing by *Mb1/3*) was greater in the AW (20.43 ± 2.59 mm g^*-1/3*^) than in the HW (13.16 ± 2.53 mm g^*-1/3*^). Wing thickness at the most proximal measured slice of the AW varied among the study species from 0.41 mm to 4.38 mm g^*-1/3*^ (mean = 2.10 mm g^*-1/3*^) and tapered to the wrist joint. Thickness of the wrist joint ranged from 0.44 mm g^*-1/3*^ to 2.36 mm g^*-1/3*^ with an average of 1.14 mm g^*-1/3*^. Wing thickness tapered further toward the most distal measured slice of the HW (range = 0.20 – 1.90 mm g^*-1/3*^, mean = 0.50 mm g^*-1/3*^). Wing cross-sectional area showed a similar pattern, tapering from a mean of 17.22 mm^2^ g^*-2/3*^ at the most proximal measured wing slice (range = 2.20 to 42.08 mm^2^ g^*-2/3*^) to a mean of 2.09 mm^2^ g^-*2/3*^ at the most distal measured slice (range = 0.64 to 19.70 mm^2^ g^*-2/3*^). The profiles of camber, chord, cross-sectional thickness (*XST*) and cross-sectional area (*XSA*) across the measured portion of the wing are shown in Fig. 2.

**Figure 2.**
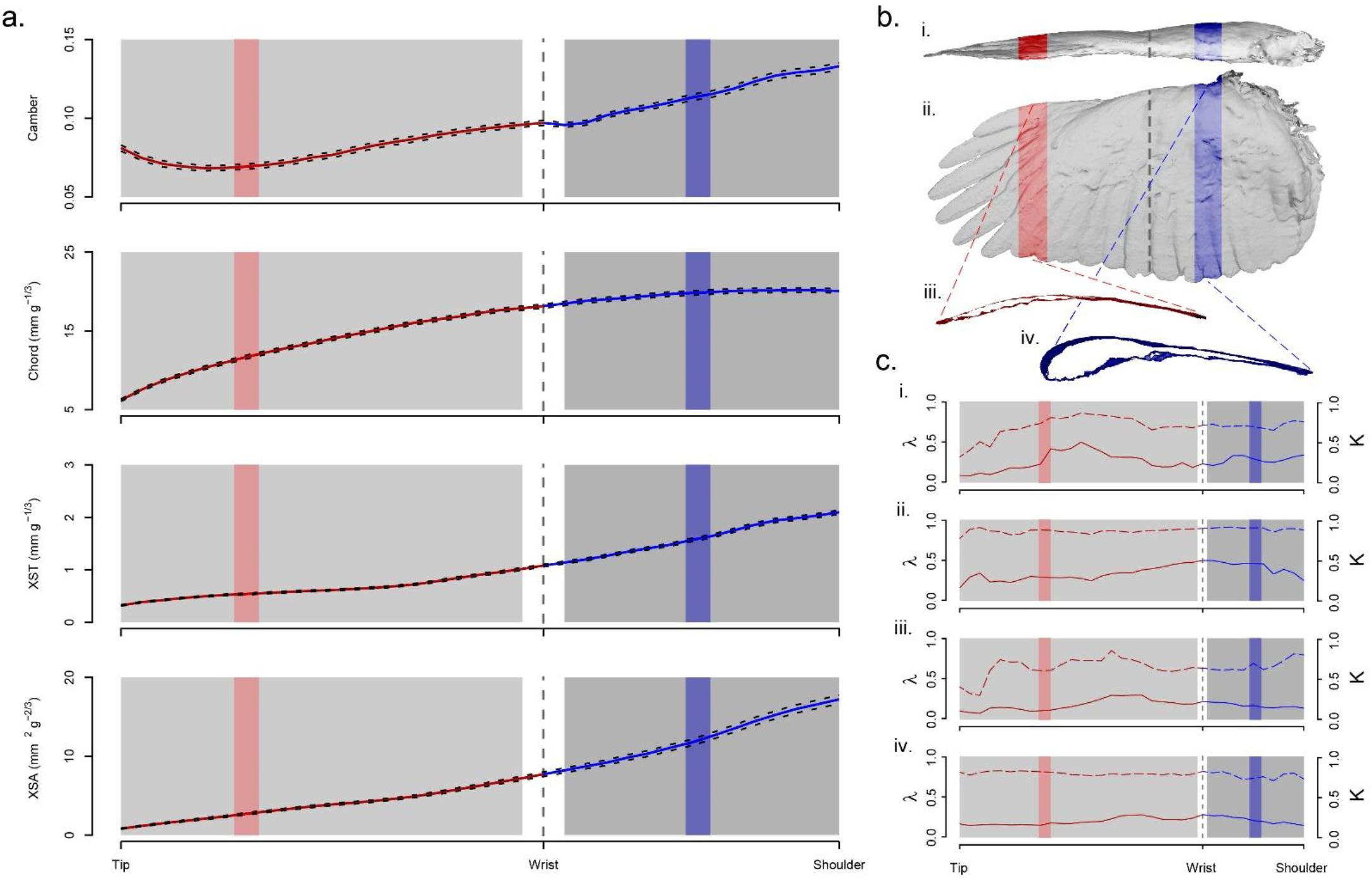
Wing shape traits scaled by body size showed marked consistency across taxa, were of greatest magnitude in the armwing (dark gray region), and decreased away from the shoulder, across the wrist joint (vertical dashed line) and through the armwing (lighter gray region). A.) Mean values (solid red/blue lines) ± SE (black dashed lines), across taxa, of camber, chord, cross-sectional thickness (*XST*) and cross-sectional area (*XSA*) in the handwing (red) and armwing (blue). The shaded gray boxes show the regions that were included in subsequent analyses of morphological and evolutionary modularity, and the red and blue shaded regions correspond with wing slices pulled from the scans (shown in Pane b.). The wrist was excluded. b.) Example of a 3D scanned wing from a Cooper’s hawk (*Accipter cooperii*) in i.) frontal and ii.) planform views, with iii.) representative slices shown for the handwing (red) and armwing (blue). c.) Phylogenetic signal was high in all shape traits in both wing regions. Dashed red/blue lines show Blomberg’s *K* for camber (i.), chord (ii.), *XST* (iii.), and *XSA* (IV.) along the length of the wing, and solid lines show Pagel’s *λ*.

### Morphological modularity

The covariance ratio (CR) test (*41*) identified significant morphological modularity (*CR* < 1.0, see Fig. 3) in the log-transformation of all shape traits (camber *CR* = 0.79, *p* < 0.001; chord *CR* = 0.79, *p* < 0.001; *XST CR* = 0.87, *p* < 0.001; *XSA CR* = 0.87, *p* < 0.001), suggesting that the AW and HW are morphologically discrete subunits of the wing. Log-transforming the data removed the biasing effect of differing means between the regions (see *42*), but a similar outcome was obtained from the raw data as well. Additionally, this result was robust to inclusion of the wrist in either the hand or arm region.

**Figure 3.**
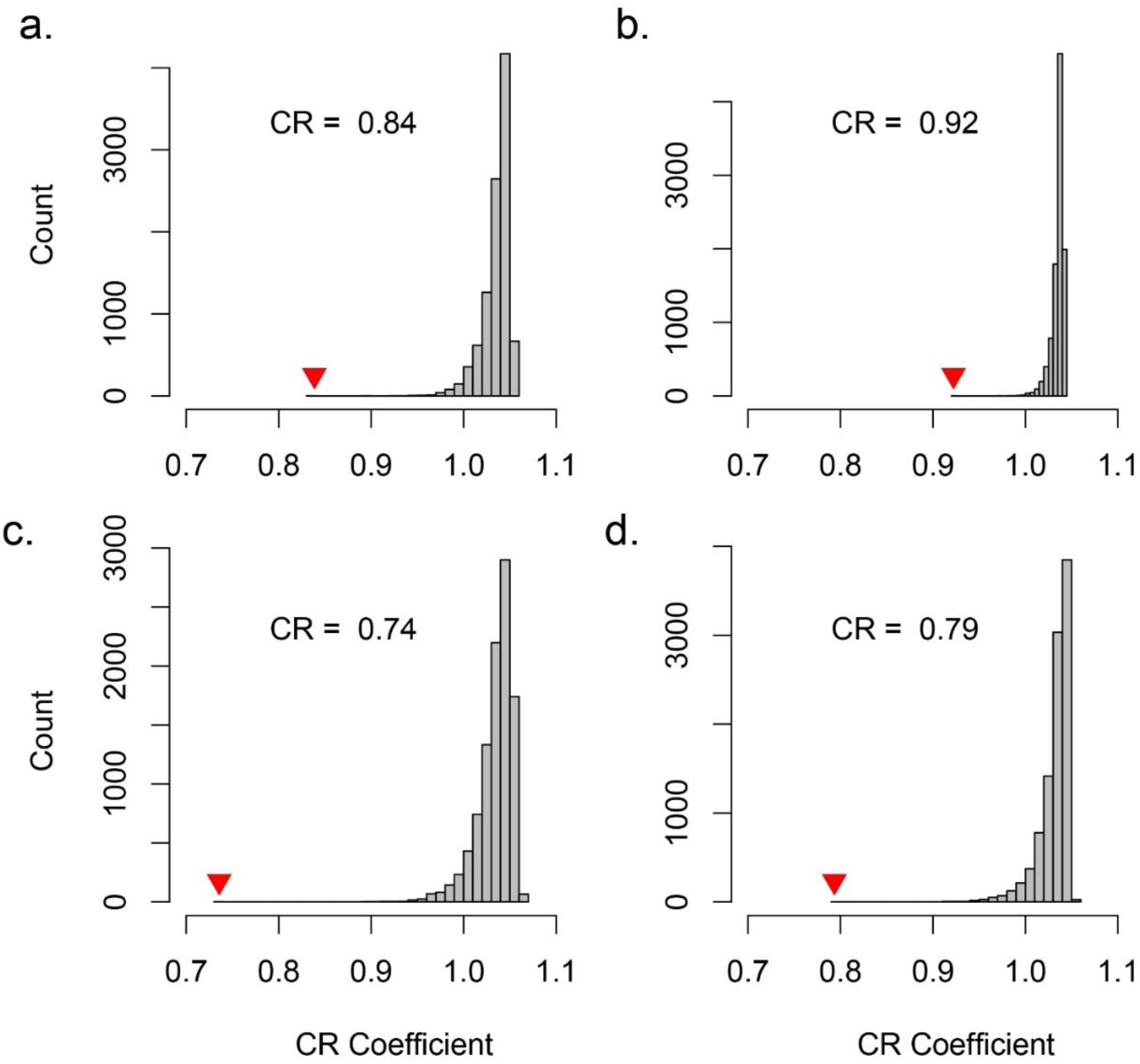
The covariance ratio test (CR) ^49^ identified significant modularity between the handwing and the armwing for a.) camber, b.) chord, c.) cross-sectional thickness, and d.) cross-sectional area. A measured CR (inverted red triangle) that is less than 1.0 indicates morphological modularity between the two wing regions. In all cases, the measured CR value was significantly less than the mean recovered from 10,000 bootstrap replicates wherein we randomly assigned slices to each of the wing regions ^49^.

### Morphological disparity

Disparity of camber was greater in the HW (mean = 0.014, see Table 1) than in the AW (mean = 0.008), with maximum (0.017) near the middle of the HW and a sharp downward transition through the wrist joint (Fig. 4). Regression-discontinuity analysis (RDA) confirmed that the spanwise trend in camber is discontinuous about the wrist (*p* < 0.001). Disparity of chord exhibited no spatial discontinuity, had comparatively low values throughout the AW (mean = 0.007), and increased distal to the wrist. Mean chord disparity in the HW was 0.012, and the distinction between the HW and the AW was supported by RDA (*p* < 0.001). Disparity of *XST* and *XSA* shared similar patterns: disparity was greatest near the tip of the wing, and decayed toward the middle of the HW (consistent with the pattern depicted in Fig. 1C, pane “ii”). In both cases, RDA showed marginal support for the discontinuity across the wrist joint (*pt* = 0.025, *pA* = 0.014).

### Evolutionary tempo and phylogenetic signal

For all shape traits, AICc supported an OU model of evolution across all wing slices (all ΔAICc > 4). Evolutionary tempo (*σ*^*2*^) among the shape traits (camber, chord, *XST*, and *XSA*) broadly showed trends similar to morphological disparity. Greater evolutionary rates corresponded with higher disparity (Table 1). However, the discontinuity across the wrist joint was less distinct, except in chord (RDA *p* < 0.001). Discontinuity results were marginal for *XST* (*p* = 0.028) and non-significant for *XSA* and camber. Phylogenetic signal was high throughout the wing in all shape traits (all K < 0.35 and all λ > 0.60, see Table 1).

## Discussion

Bird wings are complex structures composed of a feathered aerodynamic surface, skeleton, muscles, and connective tissues (Fig. 1b). The wing also contains two anatomical regions: the handwing (HW) distal to the wrist, and the armwing (AW) proximal to it. We hypothesized that this complexity results in morphological and evolutionary modularity, meaning that trait variation between anatomical subunits is greater than the variation within them, and that evolutionary dynamics differ between those modules.

We further hypothesized that the increase in mechanical sensitivity (the strength of the relationship between trait morphology and functional output (*9, 10*)) along the wing as the square of the distance from the wing base brought about by the aerodynamic force gradient and moment of inertia for flapping wings (*40*) (see Fig. 1c; Appendix 1) would result in a matching gradient in evolutionary dynamics. In earlier studies of mechanical sensitivity using 4 bar linkage systems (*9, 10*), the individual links are both modules and have discrete mechanical sensitivities, unlike the mechanical sensitivity gradient in flapping wings, and thus leave uncertain how modularity and mechanical sensitivity might interact in this case. We combined the two frameworks by hypothesizing that evolutionary dynamics would follow the mechanical sensitivity gradient, but exhibit a regionally discontinuous pattern between the HW and AW. We used a dataset of 1096 3D scanned wings from 178 bird species to assess the morphological modularity in bird wings and test whether evolutionary dynamics follow the mechanical sensitivity gradient imposed by flapping flight, the hypothesized morphological modularity, or both effects together.

We found significant discontinuities in morphological disparity across the wrist joint in all traits, supporting discretization of the HW and AW as two morphological modules. Disparity was significantly greater in the HW for all traits, and *σ*^*2*^ largely followed a similar trend. Both measures decreased away from the wingtip even within the handwing, following the gradient of mechanical sensitivity for flapping flight but breaking the strictly modular expectation for evolutionary dynamics. There was significant linear regression discontinuity for *σ*^*2*^ in wing chord and marginally for *XST*. However, *σ*^*2*^ for camber and cross-sectional area were not discretely separated at the wrist, despite greater values near the wingtip. Our results match our hypothesis that the HW and AW are morphological modules, and that evolutionary dynamics follow the mechanical sensitivity of the morphological traits. However, we also show that 1) evolutionary rate can track mechanical sensitivity within and not just among modules; thus the link between mechanical sensitivity and evolutionary rate does not depend upon morphological modularity and 2) evolutionary dynamics can track continuous mechanical sensitivity gradients.

### Bird wings are modular structures

Gatesy and Dial (*39*) proposed that locomotor modularity (the integration of anatomical subunits, such as the hindlimbs or forelimbs, into functional subunits during locomotion) is responsible for the evolutionary diversification of avian morphology and locomotion, and potentially for the origin of flight. Here, we present a refinement to their view of modularity within the structure of the wing. Our results show that bird wings are complex structures composed of at least two morphological modules, the handwing (HW) and the armwing (AW), delineated by the wrist joint. We measured morphological disparity, a quantification of the occupancy of multivariate space such as that formed by multiple morphological axes (in this case, four axes: camber, chord, *XST* and *XSA*), along the wing. We found that morphological disparity is greatest in the HW, and especially so at the wingtip. We used Regression Discontinuity Analysis (RDA) to demonstrate that the patterns of morphological disparity were different in the HW and the AW (see Fig. 1g), providing evidence that the HW and AW are discrete modules. Morphological modularity in complex biological systems is the result of shared inheritance, similar developmental patterns, or shared function among traits (*6, 7, 43, 44*). Morphological modularity has been documented in mammalian backbones, where a gradient of selective pressures along the length of the spine leads to regionalization of both form and function (*44, 45*). The flight feathers of birds’ wings form a set of serially-homologous elements akin to vertebrae in the mammalian backbone, and also experience a gradient of forces (*40*). Our results show that wing morphology shows significant regionalization, however the organization of modularity in the wing is derived from the skeleton, rather than from the feathers. Furthermore, the relationship between form and function among the wing modules is only implied and whether regionalization of morphology also leads to regionalization of biomechanical output warrants further attention.

Our finding that the HW and AW form discrete morphological modules does not imply that additional modularity cannot be found within wings. The skeletal, muscular, and integumental components of the wing might experience unique evolutionary pressures and tradeoffs that shape their evolution across multiple levels of organization (*7, 43*). For example, the thickness of the wing skeleton is tied both to its aerodynamics – thicker wings present more frontal area to the wind and produce more drag, and to its structural rigidity – thicker wing bones are more able to resist bending (*46*). The geometry of the wing’s feathered surface influences, in part, the magnitude and distribution of the forces the skeleton must withstand (*40, 47*). Therefore, these features might show morphological integration, *i*.*e*., that their morphologies coevolve within discrete regions of the wing, and the strength of the integration may vary along the length of the wing. Conversely, the flight feathers are serially homologous features arranged along the wingspan, whose relative sizes determine the dimensions and geometry of the wing surface (*17, 48*). Each feather is potentially exposed to different evolutionary pressures based on its position on the wing (*40*), raising the possibility that each feather could be its own morphological and evolutionary module. Our 3D scanned wings did not permit investigation of modularity at these levels of organization, so it possible that a greater magnitude of modularity exists that could be uncovered by different measurement techniques.

### Trait evolution follows a gradient rather than modules

We predicted and found that bird wings show strong morphological modularity between the HW and the AW. We also predicted that evolutionary dynamics (tempo and mode) would differ significantly between morphological modules. While we did find that mean values of *σ*^*2*^ were greater in the HW for all traits, evidence for evolutionary modularity was, at best, equivocal. Regression discontinuity analysis identified significant transitions in *σ*^*2*^ for chord and marginally for *XST*, but not for camber or *XSA*. Instead, *σ*^*2*^ for *XST* is consistent across much of the wing, with a notable increase near the wingtip. Camber *σ*^*2*^ also shows an increase at the wingtip but is otherwise consistent within the HW and greater than in the AW. We therefore found little support for evolutionary modularity within the wing and propose that evolutionary change of the shape traits discussed here is not beholden to morphological modules, but instead follows a smooth gradient along the span of the wing.

Aerodynamic forces in flapping flight increase as the square of the distance from the wing base (*40*), and as such, morphological alterations in wing geometry near the tip of the wing will produce outsized impacts on aerodynamic performance. The result is a span-wise gradient of increasing mechanical sensitivity toward the tip of the wing. Mechanical sensitivity influences evolutionary dynamics of morphological traits, biasing toward higher rates of evolutionary diversification (*10, 11*), as we found in the HW. Greater morphological disparity and faster evolutionary tempo in the HW (and particularly its distal tip region) relative to elsewhere along the span support our hypothesis that selective pressures driving morphological evolution in avian wings are related to the distribution of aerodynamic and inertial forces along the span of the wing. Our results are also consistent with prior work demonstrating that planform wing shape has diverged primarily near the wingtips (*17*), and demonstrate that 3D shape traits (camber, *XST*, and *XSA*) behave similarly.

Prior studies linking mechanical sensitivity to evolutionary dynamics in 4-bar linkage systems have documented transitions in evolutionary mode (i.e. Ornstein-Uhlenbeck vs. Brownian motion) in addition to a shift toward higher rates (*10, 11*). However, we found no shift in mode across the wrist joint. The OU model was best supported by AICc in all shape traits across the entirety of the wing. This is unsurprising for camber and *XSA*, as the RDA models for these traits did not highlight significant transitions in evolutionary rate (*σ*^*2*^) across the wrist joint. However, there were significant differences in *σ*^*2*^ between the wing regions for chord and, marginally, for *XST*, but without a shift in evolutionary mode. We posit that the lack of sharp transitions in evolutionary dynamics at the wrist joint, despite trait disparity analysis supporting discretization of the HW and AW into separate morphological modules, stems from the continuous gradient of increasing mechanical sensitivity along the span of the wing.

### Additional considerations

Several shape indices have been developed to facilitate the broad taxonomic sampling necessary to explore how wing shape in birds relates to their behavior and ecology. The most widely adopted of these is the handwing index (HWI) (*17, 48*), which serves as a proxy for wing aspect ratio. The present results suggest that the wingtip is evolutionarily labile and likely to be tuned to the various flight and lifestyle pressures among avian taxa, validating wingtip shape indices such as HWI. However, the utility of wingtip indices remains limited. HWI provides an imperfect proxy for wing aspect ratio. The proportion of the AW varies among avian taxa (from approximately 30 to 60% of wing length in our sample). Birds with identical HWI can have very different AR. Camber interacts with AR and plays an important role in aerodynamic force production (*33, 34*), but is not captured by any wingtip shape index. Wing volume (and by extension, mass) affects the inertial moment of the wing, influencing the energetic cost of flapping and the ability to use wing inertia for maneuvering.

We investigated the evolution of static wing shape, but bird wings are dynamic structures. Planform wing shape is dynamically and deliberately modified by birds in flight, termed “wing morphing” to modify aerodynamic performance (*49, 50*) and to react to transient perturbations (*51*–*53*). Three-dimensional shape traits like camber and span-wise twist vary as the wing cycles through its flight stroke and when acted upon by aerodynamic forces in flight (*54*), and the range of motion at the wing joints is a stronger predictor of flight style than 2D wing shape (*55*). A systematic understanding of how static wing shape affects the manner and to what degree birds can dynamically alter the shape of their wings in flight remains elusive and should prove to be a fruitful avenue for further investigation.

### Concluding remarks

We assembled an unprecedented dataset of 3-dimensional wing shape in a broad taxonomic sample of birds. Our analyses show that the wing is divided into at least two morphological modules separated by the wrist, the handwing and the armwing, and that shape divergence was greatest in the handwing. We tested competing hypotheses of how evolutionary dynamics act upon the wing modules, and found that morphological disparity was significantly modular within the wings, but that evolutionary tempo followed a gradient of mechanical sensitivity along the span of the wing that was predicted by a blade element model of flapping flight aerodynamics and inertial moment (*40*). This expands our understanding of evolutionary dynamics of complex biological structures, demonstrating that morphology can be tuned along continuous gradients in addition to previously described modular processes (*10, 11*). Our results concur with prior observations that mechanical sensitivity drives evolution of biomechanical traits. Furthermore, we demonstrate that the linkage between mechanical sensitivity and evolutionary dynamics is not specific to four-bar linkages, but also exists in other biophysical systems and therefore might be fundamental to the evolution of form and function.

## Methods

### Wing scanning and measurement

We collected three-dimensional wing shape data from spread wings in the collection at the North Carolina Museum of Natural Sciences (NCMNS) in Raleigh, NC using a NextEngine 3D Scanner Ultra HD laser scanner (NextEngine, Inc., Santa Monica, CA; Fig. 1d). Sample sizes of each taxon were limited by the availability of specimens in the NCMNS collection, but when available, we scanned 16 individuals per species, maintaining a balanced sex ratio. Scan resolution was set to optimize scanning time while preserving surface detail. Resolutions ranged from 78 dots per cm^2^ for large wings to 6300 dots per cm^2^ for smaller wings. The scanned wings (Fig. 1e) were processed using a MATLAB program (MATLAB r2014b, The MathWorks, Natick, MA, USA) that extracted the vertices from the 3D object files, creating point clouds in the shape of the wings. We used a principal components analysis to align the wing point clouds by their span (PC1, *X*-dimension), chord, (PC2, *Y*-dimension), and thickness (PC3, *Z*-dimension). Wing length (*r*) was measured as Xmax - Xmin, and wingspan as 2*r*. Because the wings were removed from the body in preservation, we were unable to account for the width of the body in our measure of wingspan.

Three-dimensional shape traits were measured by subdividing the wings into chord-wise slices along their span (Fig. 1f). To facilitate direct comparisons between wings and among taxa, we set the width of the slices to be 1/25^th^ of the distance from the wrist joint to the tip of the wing, ensuring that all wings would have the same number of handwing (HW) slices. The number of slices representing the armwing (AW) was allowed to vary, as the proportion of HW vs. AW differs among taxa. Substantial trauma occurs during the removal of the wing during preservation, so the proximal 1/3^rd^ of the AW was excluded from analyses to reduce the influence of preservational artifacts.

We measured four shape traits from each wing slice. Chord was measured as *Y*_*max*_ - *Y*_*min*_, and cross-sectional area (*XSA*) was measured as the area contained within a spline fitted to the perimeter of the wing section in the *Y / Z* plane. The maximum distance in the Z-dimension (i.e., the greatest distance from the upper wing surface to the lower wing surface) was recorded as the maximum cross-sectional thickness (*XST*). Camber was calculated as (*Z*_*max*_ - *Z*_*min*_) / (*Y*_*max*_ - *Y*_*min*_).

Body mass (*Mb*) was recorded from museum tag data where available. When mass was not available from specimen tags, a species mean value was filled in from the CRC Handbook of Avian Body Masses (*56*). Measurements of wing length, area, chord, and thickness were scaled by dividing by body mass taken to the appropriate power (*Mb*1/3 for linear measures and *Mb*2/3 for areas) and summarized within each taxon. Subsequent analyses were conducted on species median values for each wing slice.

### Phylogenetics

Phylogenetic analyses were based upon the Jetz. *et al*. (*57*) super tree from Birdtree.org (*58*). The tree was pruned to include only taxa in our scanned wing dataset. Handling of the tree, data, and phylogenetic analyses was done using tools from the *Phytools* (*59*) and *geiger* (*60*) packages in the R Statistical Computing Environment version 4.1.0 (*61*). Phylogenetic signal (i.e. Blomberg’s *K* (*62*) and Pagel’s *λ* (*63*)) was calculated for each wing slice using the ‘*phylosig*’ function in the *Phytools* package.

### Morphological modularity analysis

#### Modularity

To assess whether the HW and AW are morphologically distinct modules, we used the covariance ratio (*CR*) proposed by Adams (*41*). This test compares the covariance among traits within a putative module to covariance among the modules. The test statistic (*CR*) ranges from 0 to positive infinity, with values between 0 and 1 representing greater covariance within putative modules than among them, signaling morphological modularity. *CR* greater than 1 indicates morphological integration, and a lack of modularity (*41*). This test was implemented using code provided in the supplement of Adams’ description of the method (*41*). Because the mean values of the shape traits differ between wing regions, in addition to conducting the modularity test on the isometrically scaled data, we used a log10-transformation to mitigate any biasing effect from the difference in means. Furthermore, because the wrist joint affects wing camber, thickness, and *XSA*, its effect on the modularity analysis was difficult to predict. To test whether inclusion of the wrist influenced our modularity interpretations, we iterated our analysis, including the wrist in each the HW and the AW regions while excluding it from the other. We also removed the wrist from consideration entirely, and only analyzed regions proximal and distal to it. Because inclusion of the wrist had no impact on the modularity analysis, and for the sake of simplicity, we present results excluding the wrist since this avoids arbitrarily assigning it to either the AW or HW region.

#### Disparity

Morphological disparity is a measure of the variation in traits among taxa. We compared morphological disparity in each of our shape traits for each wing slice using the ‘*dispRity’* function in the *dispRity* package (*64*) in R. We compared disparity between the HW and the AW using regression discontinuity analysis (RDA) using the *‘rdd_reg_lm’* function in the *rdd_tools* package (*65*) in R. Regression discontinuity analysis is a statistical tool to assess changes in slope or intercept in a temporal or spatial trend across an assigned *X*-axis cutoff point, in our case, the wrist joint. A difference in disparity in the same shape traits between different regions of the wing would indicate a difference in evolutionary lability as well.

### Evolutionary modularity analysis

#### Evolutionary tempo and mode

To test whether different morphological modules expressed different evolutionary dynamics, we fit Brownian motion (BM), Ornstein-Uhlenbeck (OU), and early-burst (EB) evolutionary models to each of the shape traits at each of the wing slices using the ‘*fitContinuous*’ function in the *geiger* package in R. We used Akaike’s Information Criterion corrected for small sample size (AICc) to determine the most suitable model for each trait and estimated evolutionary rate (*σ*^*2*^) for each wing slice using that model. We tested for differences in patterns of *σ*^*2*^ between the wing regions using regression discontinuity analysis (RDA) as described above.

## Acknowledgements

We thank the North Carolina Museum of Natural Sciences, especially Brian O’Shea and John Gerwin, for access to specimens and assistance throughout the wing scanning operation. We also thank Alaowei Amanah, Eli Bradley, Sophia Chizhikova, Elliot Cho, Lucy Herrero, Russel Lo, Raghu Padma, Alva Rönn, Colton Sanders, Sarah Yaghoubi, and Zhitong Yu, for their tireless efforts scanning wings. Pranav Khandelwal provided thoughtful discussion and feedback throughout the project. Lindsay Waldrop, P. Khandelwal, Brenna Hansen and Sonja Friman gave helpful feedback on an earlier draft of the manuscript.

## Funding

National Science Foundation (IOS-1253276) to TLH,

NSF DEB 1737752 to Daniel R. Matute,

Sigma Xi for a GIAR award to JAR, and

NextEngine, Inc. for providing financial support for wing scanning equipment and software.

## Author contributions

Conceptualization: JAR, TLH

Methodology: JAR, TLH

Investigation: JAR, TLH

Visualization: JAR

Supervision: TLH

Writing—original draft: JAR

Writing—review & editing: JAR, TLH

## Supplementary Materials for

## Supplementary Text

### Mechanical sensitivity models for flight

Here we elaborate on the underlying mechanical sensitivity models for the wings of flying animals described in the main text. Mechanical sensitivity relates the rate of change in some aspect of morphology to the rate of change in a relevant biomechanical output. Here we select aerodynamic lift as the most relevant biomechanical parameter for flight. All flying animals must generate lift sufficient to support their body weight, whereas other potentially relevant aerodynamic parameters such as drag and the mechanical power required for flight are not similarly constrained, making them less than ideal candidates for these models. However, variation in overall body size means that the total amount of lift differs among birds, so a general model of mechanical sensitivity must be normalized as a percent change in lift for a percent change in morphological properties.

### Flapping

We model mechanical sensitivity in flapping flight following the strips (i.e. blade-element) analysis popularized by Weiss-Fogh ^40^. In brief, this analysis divides the wing into a set of infinitesimally thin chord-wise strips and integrates the lift produced by each strip along the root to tip length of the wing. The lift produced by an airfoil is calculated as:

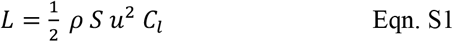

where *L* is lift, *S* is airfoil area, *u* is flow velocity past the airfoil, and *C*_*l*_ is the coefficient of lift, a non-dimensional quantity. We use a subscript *i* to denote the lift from the *i*th section, and express area *S* as the produce of the section width *dr* and section chord length *c*_*i*_.

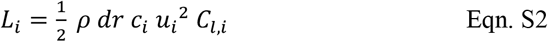

Equation S2 provides the lift for each strip in terms of the flow velocity *u*_*i*_ past the strip. For flapping flight, this flow velocity also varies from base to tip due to the motion of the wing. For hovering flight, velocity is zero at the base, reaches a maximum at the tip, and is given by:

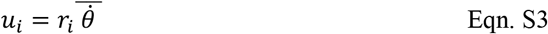

where 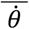 is the average angular velocity of flapping. Substitution, of Eqn. S3 into Eqn. S2 produces:

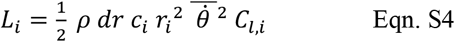

This result can then be integrated across the wing radius:

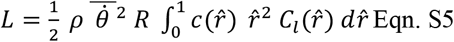

where 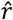 is the non-dimensional radius, i.e. the actual radius *r*_*i*_ of the strip divided by the length of the wing, *R*. Equations S4 and S5 show that for a wing of length *R*, the percent change in lift at a particular spanwise section *i* or non-dimensional spanwise location 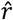 produced by a change in chord or coefficient of lift depends on 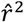. For example, increasing *C*_*l*_ by 20% at 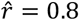 versus 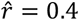 produces 0.8^2^0/.4^2^.=4times the increase in lift, thereforeis has 4 times greater mechanical sensitivity. This produces the following curves:

**Figure S1:**
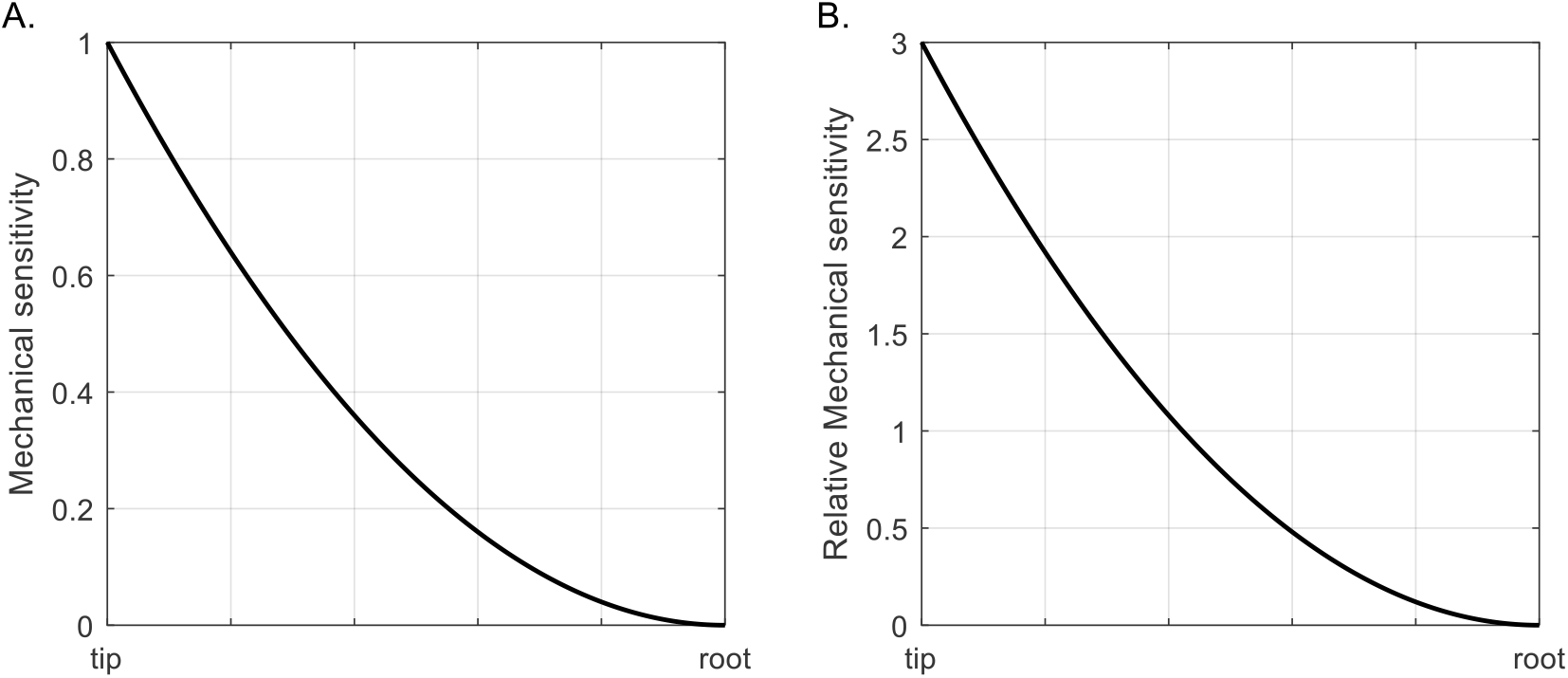
mechanical sensitivity in flapping. These curves show the predicted mechanical sensitivity of wing sections from root to tip. Panel A is strictly in terms of 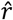 while panel B has the result multiplied by three so that the average value of the curve is 1.0 to simplify comparison with gliding wings.

Figure S1 directly predicts the mechanical sensitivity for quantities such as chord that have a direct, linear relationship with lift in Eqn. 5. However, of the four quantities we measured: camber, chord, cross-sectional area, and cross-sectional thickness, only chord meets this criterion. The other three parameters: camber, cross-sectional area, and cross-sectional thickness are all expected to affect the coefficient of lift *C*_*l*_, but may not have a simple linear relationship with it. However, assuming that the relationship, whatever it is, is similar among all wing sections, the underlying mechanical sensitivity curve will be as shown in Fig. S1.

### Gliding

In gliding flight all wing sections have the same velocity with respect to the flow, such that the above analysis for flapping flight produces a flat prediction of mechanical sensitivity, with no differentiation along the wing. In practice, idealized models of lift production from fixed wings produce and elliptical distribution with lift reaching zero at the tip. However, the details of this idealized elliptical distribution vary with wing shape, with greater sensitivity toward the tip where the geometry may affect the shape and magnitude of the tip vortex. In the absence of an analytic expression for these interactions, we hypothesize no variation in mechanical sensitivity for gliding flight along the span of the wing as shown in Fig. 1c.

### Inertia

When birds flap their wings, they use mechanical power to overcome the inertia of the wing, with the cost depending on the moment of inertia of the wing as well as flapping frequency and amplitude ^35^. Thus, wing moment of inertia might also produce a mechanical sensitivity gradient in the wings of flying animals. The moment of inertia for a wing flapping about its base is given by

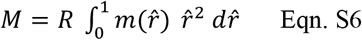

where 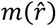 is the mass of the wing at non-dimensional spanwise location 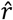 Note that this equation produces the same general mechanical sensitivity curve that is created by the aerodynamics of flapping (Fig. S1), but depends on wing mass instead of wing chord or coefficient of lift. Thus, considering inertia as well as lift does not alter the overall mechanical sensitivity of flapping flight.

